# Salidroside Mitigates Malignant Arrhythmias by Restoring Sodium Channel Function During Ultra-Acute Myocardial Infarction

**DOI:** 10.1101/2024.07.31.606101

**Authors:** Gongxin Wang, Yilin Zhao, Chenchen Zhang, Xiuming Dong, Siyu Sun, Xiulong Wang, Dongxu Li, Xuefang Li, Huan Li, Chieh-Ju Lu, Yimei Du, Zhigang Chen, Fei Lin, Guoliang Hao

## Abstract

**Background:** The ultra-acute phase (Phase 1a) of acute myocardial infarction (AMI) is marked by a high incidence of malignant arrhythmias, often occurring during the prehospital period. Currently, there are no effective treatment options available for managing these arrhythmias at this early stage.

**Methods and Results:** Using dual-channel optical mapping, we simultaneously recorded membrane potentials and calcium transients during acute myocardial infarction. Calcium transient duration maps accurately localized the infarcted region, and action potential activation time maps revealed conduction heterogeneity in the infarcted zone. Patch-clamp recordings showed that Salidroside (Sal) (1 µg/mL) significantly increased sodium current density from -59.27 ± 2.15 pA/pF to -83.46 ± 3.19 pA/pF (P<0.01) and shifted the Nav1.5 activation curve leftward (V1/2 from -37.27 ± 0.5 mV to -44.55 ± 0.7 mV, P<0.01). In rat and rabbit AMI models, Sal pre-treatment reduced conduction heterogeneity and arrhythmia incidence compared to controls. Optical mapping showed improved conduction velocity and uniformity in the Sal group.

**Conclusions:** Sal restores electrophysiological function in damaged myocardium by modulating sodium currents, reducing conduction heterogeneity, and decreasing malignant arrhythmia incidence during the ultra-acute phase of AMI. These findings suggest a novel therapeutic strategy for AMI, addressing a critical unmet need in antiarrhythmic therapy.

**What is New?:**

1. This study identifies Salidroside (Sal) as a novel agent that enhances sodium channel currents (Nav1.5), distinguishing it from traditional antiarrhythmic drugs which primarily target potassium channels or β-adrenergic receptors. Sal improves conduction uniformity in the infarcted myocardium by restoring the electrophysiological function of damaged cardiac cells, eliminating slow conduction pathways, and reducing conduction heterogeneity.
2. This research introduces Sal as a promising candidate for preventing and treating arrhythmias during the critical early stages of MI, potentially improving patient outcomes. Sal administration during the ultra-acute phase (phase 1a) of myocardial infarction (MI) significantly reduces the incidence of malignant arrhythmias, a critical period characterized by high extracellular potassium and increased arrhythmia risk.
3. Utilizing calcium transient imaging and optical mapping, this study provides precise localization of ischemic regions and detailed electrophysiological characterization, offering a robust methodology for assessing therapeutic efficacy.

## Introduction

Acute myocardial infarction (AMI) remains a leading cause of sudden cardiac death globally, posing a significant challenge to cardiovascular medicine^1–3^. The rapid progression of AMI, especially during the ultra-acute phase (phase 1a), often leads to malignant arrhythmias, which are a major contributor to mortality^4, 5^. Phase 1a occurs approximately 6-15 minutes after the onset of AMI and frequently precedes the arrival of prehospital medical care^6, 7^. Currently, there are no effective treatment options available during this critical period^2, 8^. Therefore, the prevention or treatment of malignant arrhythmias in Phase 1a is particularly important for individuals at high risk of AMI^9^.

Accurately identifying critical ischemic sites is essential for studying the electrophysiological changes that occur in myocardial ischemia following an infarction. In constructing a myocardial infarction model through ligation of the left anterior descending (LAD) artery, despite efforts to ensure consistent ligation placement, variability in vascular distribution can result in differences in ischemic regions. Due to the minimal extent of myocardial necrosis at this early stage, histopathological examinations often fail to confirm the presence of myocardial infarction^10^. Calcium transient regulation in myocardial cells involves two phases: the upstroke phase (Ton) and the downstroke phase (Toff). The Ton phase is mediated by L-type calcium channels and ryanodine receptors (RYR2), while the Toff phase is regulated by the Na+/Ca2+ exchanger (NCX), calcium pumps, and sarcoplasmic reticulum Ca2+-ATPase (SERCA), all of which are energy-dependent processes^11–13^. Consequently, we hypothesize that the duration of the transient (Toff) may be sensitive to changes induced by myocardial infarction. To test this hypothesis, we employed dual-channel optical mapping^14^ to assess calcium transient alterations during the ultra-acute phase of myocardial infarction.

Ventricular tachycardia and fibrillation can arise from various arrhythmogenic mechanisms, which may act either independently or in combination^15^. Key pathophysiological conditions associated with this phase include elevated extracellular potassium (K+), acidosis, and anoxia^16^. Despite this, the specific factors that influence an individual’s susceptibility to arrhythmias associated with ischemia are not yet fully understood. Elevated [K+]_o_ plays a critical role in determining conduction patterns, causing supernormal conduction, depressed conduction, and conduction block as [K+] levels rise from 4.5 to 14.4 mmol/L^17^. While delayed conduction has been widely recognized, the heterogeneity of conduction within the infarcted region can now be vividly illustrated through advanced optical mapping techniques.

Rhodiola is the dried root and rhizome of Rhodiola crenulata (HK. f. et.Thoms) H.Ohba, it has been used in Europe and Asia for a long time to mitigate altitude sickness.^18–20^. Altitude sickness occurs because the body struggles to adapt to lower oxygen levels at high altitudes, leading to hypoxia^20, 21^. Similarly, in the early stages of myocardial ischemia, myocardial cells face the challenge of hypoxia due to insufficient blood supply. Therefore, it can be hypothesized that Rhodiola has a similar protective effect on myocardial cells under hypoxic conditions caused by myocardial ischemia. There are 282 kinds of chemical components in Rhodiola, and the representative active component is salidroside (Sal), which has anti-hypoxia, anti-oxidative stress, anti-inflammation, anti-apoptosis, and other effects^22–25^.

This study investigates the efficacy of Sal in addressing these challenges. We hypothesize that Sal’s modulation of sodium currents could mitigate the arrhythmogenic effects of high potassium environments and improve conduction in ischemic myocardial tissue. Our findings suggest that Sal not only improves conduction velocity and reduces dispersion but also offers a novel approach to preventing malignant arrhythmias during the critical ultra-acute phase of AMI. By detailing changes in calcium transients and using these alterations to pinpoint ischemic areas, we provide a comprehensive understanding of electrophysiological disturbances and the therapeutic potential of Sal in AMI.

## Materials and Methods

### Data Availability

Supporting data, analytical methods, and study materials relevant to this research can be accessed upon reasonable request. Detailed experimental procedures, including dual-channel optical mapping, pacing protocols, electrical mapping, whole-cell patch clamping, as well as other molecular and statistical techniques, are thoroughly described in the Supplementary Data.

### Ethics and Study Approval

All procedures were approved by the Institutional Animals Ethics Committees at Henan SCOPE Research Institute of Electrophysiology Co. Ltd. (approval reference number LLSCOPE2023-121) and conducted in accordance with national guidelines. New Zealand white rabbits (male, 2.5-3.5 kg) and Sprague-Dawley rats (male, 200-250 g) were supplied by the Experimental Animal Center of the Medical College of Xi’ an Jiaotong University and SPF (Beijing) Biotechnology Co. Ltd., respectively. Animals were housed in a pathogen-free facility and sacrificed humanely via intraperitoneal injection of pentobarbital sodium (50 mg/kg).

### Pharmacological Treatment

Salidroside (Sal, CAS No.: B20504, Shanghai Yuanye Biotechnology Co., LTD) was dissolved in physiological saline. For pre-protection studies, Sal was administered to rats and rabbits, with acute studies involving perfusion at 1, 5, and 25 µg/ml concentrations, typically 15 minutes before myocardial infarction (MI) induction.

### Langendorff-Perfused Isolated Hearts

Hearts from euthanized SD rats and rabbits were excised and mounted on a Langendorff system. They were perfused with modified Tyrode’s solution at 37°C for 15 minutes before experimental procedures.

### Myocardial Infarction Model

MI was induced by occlusion of the left anterior descending (LAD) coronary artery using a 6-0 silk suture. Sham surgeries were performed as controls.

### Acute Isolation of Cardiomyocytes

After Langendorff perfusion, hearts were perfused with an enzyme solution (collagenase Type II, hyaluronidase, CaCl2) for 15-20 minutes, then ventricles were minced and agitated. The cell suspension was filtered, centrifuged, and resuspended in KB solution. Calcium was reintroduced, and cell viability was assessed.

### Whole-cell Patch Clamping

For INav1.5 and AP recording, pipette and bath solutions were used as specified. Data were collected using an Axopatch 700A amplifier, Digidata 1440 interface, and pClamp 10.6 software, with recordings conducted at room temperature.

### Optical Mapping

Stabilized Langendorff-perfused hearts were pre-perfused with pluronic F127 and dye-loaded with Rh237 and Rhod2-AM. Fluorescence was captured using a system with 530 nm LEDs, filtered for voltage and calcium signals, and synchronized using OMapRecord 5.0 software.

### Electrical Mapping

Field potentials from the epicardium were recorded with a 64-channel MEA system. Epicardial activation maps and conduction velocity were assessed using EMapScope 5.0 software.

### Stimuli Pacing Protocol

Pacing was performed with a constant voltage/current stimulator, using an amplitude of 1.5 times the diastolic threshold and a 2 ms pulse width. A pacing protocol of 6 Hz (rats) and 3 Hz (rabbits) was applied before and after MI induction or drug administration.

### Statistical Analysis

Data were analyzed with Clampfit 10.6, OriginPro 8.0, and Adobe Illustrator 10. Activation and inactivation curves were fitted with the Boltzmann equation. Statistical significance was determined by one-way ANOVA followed by multiple comparisons, with *p*< 0.05 considered significant.

## Results

### Electrophysiological Changes and Calcium Transients Post-Myocardial Infarction

We used dual-channel optical mapping to simultaneously record action potential signals and calcium transient signals from Langendorff-perfused hearts (Fig. 1A). In the ultra-acute phase of myocardial infarction, we observed that the duration of calcium transients was prolonged within 1 minute after ligation (Fig. 1B).

**Figure 1.**
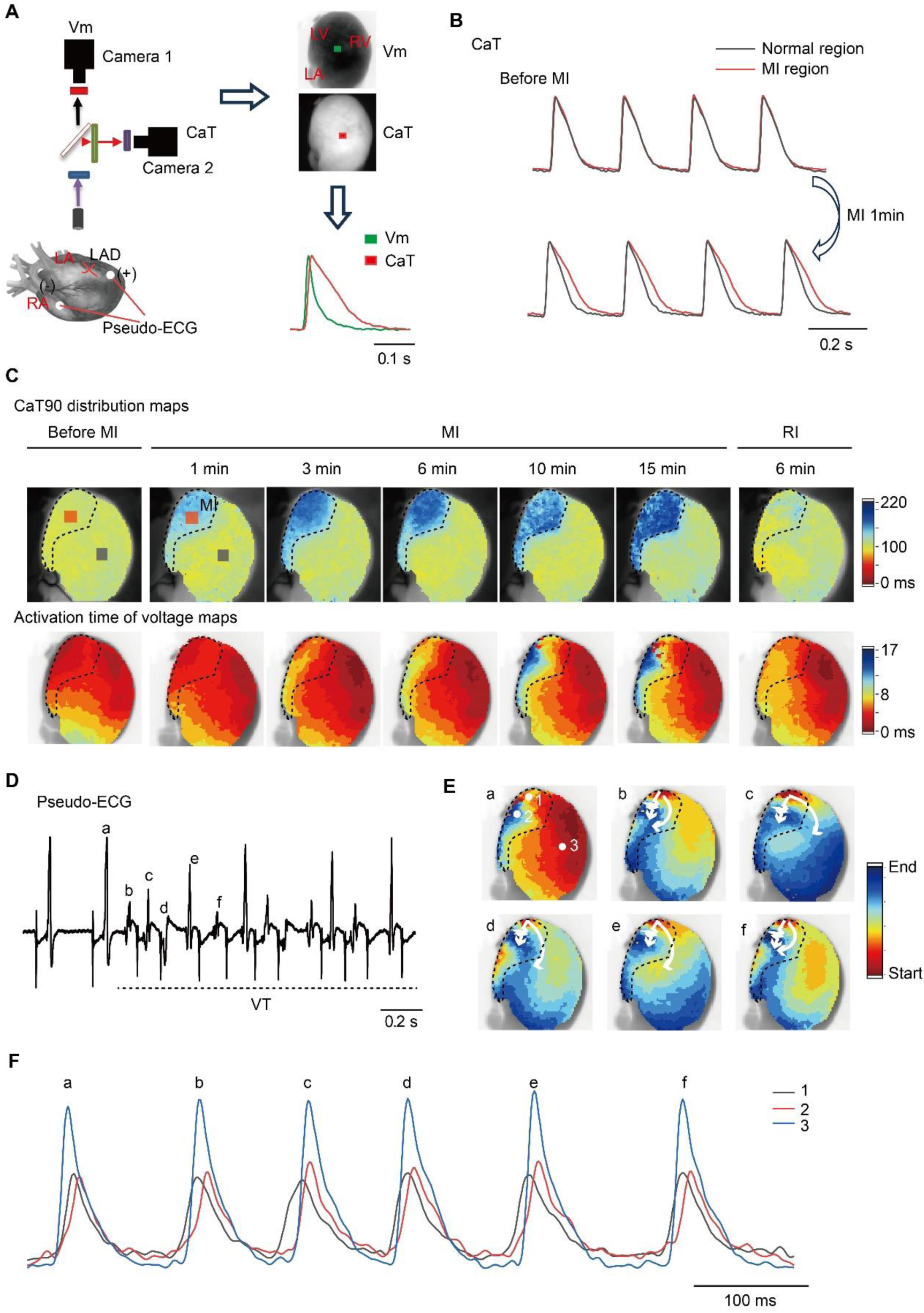
Electrocardiographic and Calcium Transient Changes Post-Myocardial Infarction. (A) Diagram illustrating the construction of the myocardial infarction (MI) model through left anterior descending artery ligation. Despite standardization efforts, variability in ischemic regions was observed. (B) Calcium transient duration extended within 1 minute of ligation, indicating ischemic areas. CaT90 distribution shows prolonged calcium transients in the infarcted region. (C) Action potential activation times before and at 15 minutes post-MI, with conduction delays evident in infarcted areas and partial normalization after reperfusion. (D) Representative ventricular tachycardia (VT) episode recorded with Pseudonym-ECG during the ultra-acute phase. (E) Conduction patterns in infarcted areas reveal both slow and fast conduction regions. (F) Action potentials at different sites (1, 2, and 3) show reduced amplitudes and earlier activation in infarcted regions compared to normal areas.

Fig. 1C shows the distribution of CaT90 (calcium transient duration) before myocardial infarction, 15 minutes after infarction, and following reperfusion. The regions with prolonged calcium transients correspond to the ischemic areas (MI regions). Fig. 1C, second column, depicts the isochronal maps of action potential activation before and 15 minutes after myocardial infarction. These images visually illustrate the progressive conduction delay in the infarcted area, which can be alleviated following reperfusion. During the 6–15-minute window, which is a critical period for the high incidence of malignant arrhythmias (phase 1a of the ultra-acute phase), we observed not only areas of conduction delay but also regions of premature excitation within the infarcted zone. Fig. 1D presents a representative example of ventricular tachycardia (VT) recorded during phase 1a in the pseudo-ECG. Fig. 1E analyzes the conduction maps of this VT episode, showing regions of slow and fast conduction within the infarcted area. The fast-slow conduction pattern is a key basis for the development of malignant arrhythmias. Fig. 1F shows action potential waveforms from three sites marked with white circles in Fig. 1E. Points 1 and 2 are located within the infarcted region, with Point 1 near an ectopic pacing site and Point 2 in a slow conduction zone, while Point 3 is in a normal region. Notably, Points 1 and 2 exhibit reduced action potential amplitudes compared to Point 3, with Point 1 being the site of earliest excitation.

### Impact of Hyperkalemia on Cardiac Myocyte Excitability

To investigate cellular excitability, we modified our protocol for eliciting action potentials by using a brief, high-current injection (e.g., 1 nA, 2 ms) to a more gradual, low-current injection (e.g., 130 nA, 50 ms) (Fig. 2A). This adjustment allowed us to precisely assess myocardial cell excitability by gradually reaching the threshold potential, as shown in Fig. 2B. This approach provides a more accurate measure of excitability based on the duration of the current injection required to induce an action potential. An increase in cellular excitability would be indicated by a reduced current needed to trigger an action potential, reflecting as a leftward shift in the plot. Conversely, a decrease in excitability would be indicated by a larger current required, showing as a rightward shift in the plot. Although prolonged low-current injections can affect the repolarization morphology, they do not impact the determination of depolarization.

**Figure 2.**
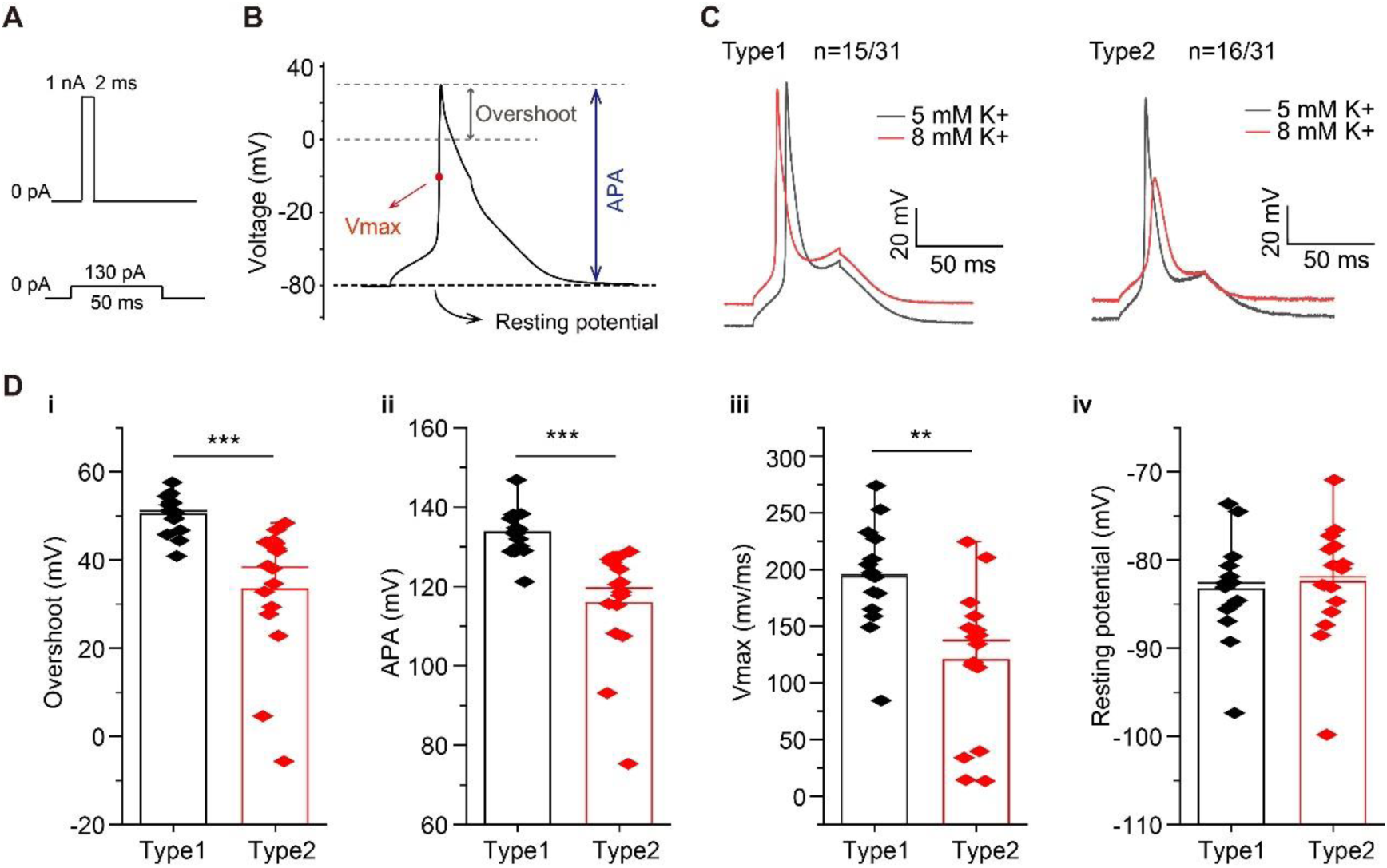
Impact of Hyperkalemia on Cardiac Myocyte Excitability. (A) Experimental setup for assessing excitability through high-current (1 nA, 2 ms) and slow-current (130 nA, 50 ms) stimulation. (B) Shift in excitability thresholds with increased excitability moving leftward and decreased excitability moving rightward. (C) Two distinct responses to 8 mM potassium: Type 1 (n=15) with increased excitability and Type 2 (n=16) with decreased excitability. Type 1 cells exhibit higher overshoot, APA, and Vmax compared to Type 2 cells. Resting potential remains unchanged. *p<0.05, **p<0.01, ***p<0.001.

Following myocardial infarction, the extracellular environment exhibits elevated potassium levels. While literature reports indicate potassium concentrations exceeding 10 mM, we chose to validate our findings at 8 mM due to its significant impact on excitability. Our experiments revealed that adult rat myocardial cells isolated via Langendorff perfusion exhibited two distinct responses to the 8 mM high-potassium environment: Type 1 and Type 2 (Fig. 2C). We analyzed the action potential parameters of both cell types before exposure to high potassium, including action potential amplitude (APA), overshoot, maximum depolarization rate (Vmax), and resting potential. Statistical analysis revealed that Type 1 cells exhibited higher overshoot, greater APA, and larger Vmax compared to Type 2 cells, with no significant difference in resting potential between the two types. This suggests that electrophysiologically normal myocardial cells are more likely to exhibit Type 1 behavior, indicating increased excitability, in a high-potassium environment. In contrast, cells with electrophysiological deterioration are more likely to exhibit Type 2 responses, indicating reduced excitability, under high potassium conditions.

### Sal Modulates Sodium Currents in Cardiac Myocytes

We recorded sodium currents in acutely isolated myocardial cells using a patch-clamp technique. The stimulation protocol involved ramp voltage stimulation, where the membrane potential was clamped at -80 mV and then ramped to +60 mV before returning to -80 mV. The depolarizing ramp stimulation induced inward sodium currents. After perfusion with 1 µg/ml of Sal, we observed that sodium currents exhibited earlier activation and increased magnitude. Figure 3A illustrates the time course of sodium current recordings. The effect of 1 µg/ml Sal on enhancing inward sodium currents was rapid, becoming evident within approximately 100 seconds, and the effect could be washed out and re-induced upon reapplication. Figure 3B presents a statistical analysis of the effect of 1 µg/ml Sal on Nav1.5 current density. Sal significantly increased the current density from -59.27 pA/pF to -83.46 pA/pF, with a subsequent washout bringing it back to -59.13 pA/pF.

**Figure 3.**
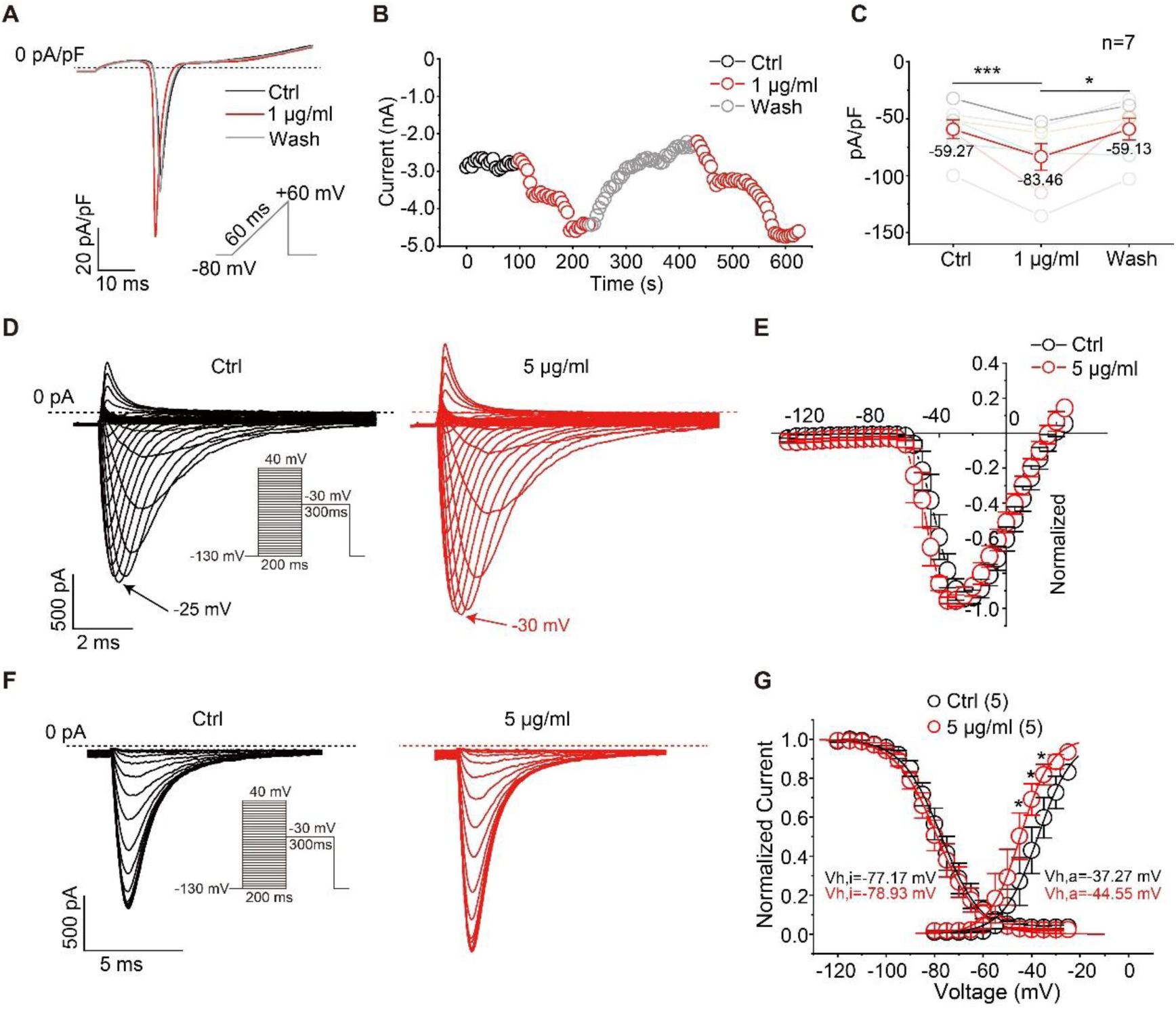
Sal Enhances Sodium Currents in Cardiac Myocytes. (A) Patch-clamp recordings show Sal (1 µg/ml) results in earlier activation and increased sodium current density. (B) Current-time plot demonstrating rapid onset and washout effects of Sal on sodium currents. (C) Increased Nav1.5 current density from -59.27 pA/pF to -83.46 pA/pF following Sal treatment. (D) Step voltage stimulation confirms Sal shifts activation curve leftward, with minimal effect on inactivation curve (E-G). *p<0.05, **p<0.01, ***p<0.001.

To further validate the effect of Sal on the Nav1.5 channel, we examined its impact on a cell line stably expressing Nav1.5 using step voltage stimulation. The voltage was clamped at - 130 mV, and then stepped from -130 mV to +40 mV in 5 mV increments for 200 ms, followed by recording the inactivation current at -30 mV. Figure 3E shows the I-V curves for control and cells perfused with 5 µg/ml Sal. Figure 3G displays the activation and inactivation curves before and after Sal administration. Notably, 5 µg/ml Sal shifted the activation curve to the left, with the half-maximal activation voltage shifting from -37.27 mV to -44.55 mV. In contrast, the inactivation curve remained largely unchanged, with the half-maximal inactivation voltage shifting from -77.17 mV to -78.93 mV.

### Restoration of Electrophysiological Properties by Sal

Action potentials in acutely isolated myocardial cells were recorded using a patch-clamp technique in current-clamp mode. Initially, we evoked action potentials by applying a 2 ms, 1 nA current injection. Subsequently, the cells were exposed to different concentrations of salidroside (Sal): 1 µg/ml, 5 µg/ml, and 25 µg/ml. We observed that Sal administration led to earlier depolarization of the action potentials following stimulation, with a noticeable increase in overshoot (Fig. 4A).

**Figure 4.**
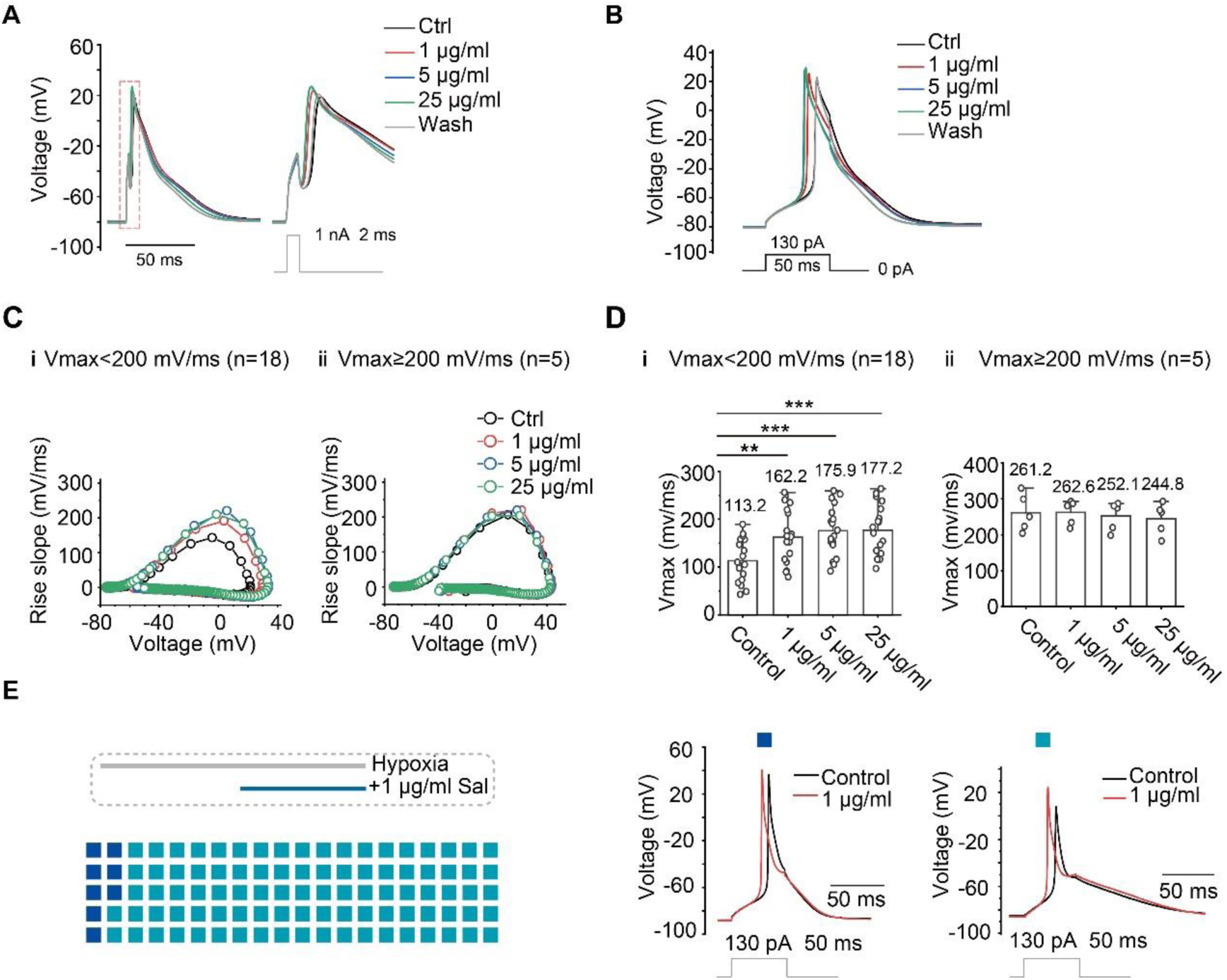
Restoration of Electrophysiological Properties by Sal. (A) Action potentials recorded from acutely isolated myocardial cells with varying Sal concentrations (1 µg/ml, 5 µg/ml, and 25 µg/ml). Sal treatment leads to earlier depolarization and increased overshoot. (B) Action potentials elicited with small, long-duration current injections, showing increased cellular excitability with Sal administration. (C) Relationship between voltage change and rate of voltage change, demonstrating increased maximum depolarization rate (Vmax) and overshoot in electrically impaired myocardium with Sal treatment. (D) Statistical changes in Vmax for myocardial cells with normal and impaired electrical function before and after Sal treatment. (E) Effects of Sal on action potential activation in hypoxic myocardial cells, with 23 of 25 cells showing earlier activation and enhanced overshoot. *p<0.05, **p<0.01, ***p<0.001.

To further investigate the effects of Sal on myocardial cell excitability, we employed a protocol involving small, long-duration current injections. This approach allowed us to achieve gradual depolarization of the membrane potential, and action potentials were elicited when the threshold voltage was reached. After applying different concentrations of Sal, we noted that action potentials were elicited with smaller amounts of injected charge, indicating an increase in cellular excitability (Fig. 4B).

Figure 4C presents the relationship between the voltage change and the rate of voltage change at different voltages. This analysis provides a clearer view of changes in the maximum depolarization rate and overshoot amplitude. We found that Sal increased the maximum depolarization rate (Vmax) and overshoot in electrically impaired myocardium (Vmax < 200 mV/ms). However, there were no significant effects on myocardial cells with normal electrical function (Vmax > 200 mV/ms). Figure 4D shows the statistical changes in Vmax for the two conditions. Additionally, figure 4E demonstrates the effects of Sal on action potential activation in hypoxic acutely isolated myocardial cells. We recorded from 25 cells, of which 23 exhibited earlier activation and enhanced overshoot after Sal treatment. Two cells showed earlier activation but did not exhibit a significant increase in overshoot.

### Optical Mapping of Salidroside Effects on the Ultra-Acute Phase of Myocardial Infarction

Figure 5A illustrates the electrophysiological changes in the isolated rat heart MI model before and after LAD ligation. Optical mapping was performed every minute for 8 seconds, with continuous recordings for 15 minutes. The figure displays the electrophysiological states before MI and at 3-, 8-, and 15-minutes post-MI. The first row presents activation time isochrones. Under sinusoidal conditions, the optical mapping of the epicardial surface shows a shorter activation time with uniform conduction. At 3 minutes post-MI, localized conduction delays are observed; at 8 minutes, larger areas of conduction delay emerge, and by 15 minutes, significant differences in excitation within the infarcted region are apparent. The second row shows the distribution of action potential (AP) amplitude values. Prior to MI, the amplitudes are uniform; however, as the duration of infarction increases, the AP amplitudes in the infarcted region decrease compared to normal areas. The third row provides the distribution of calcium transient durations, clearly depicting the ischemic regions following myocardial infarction.

**Figure 5.**
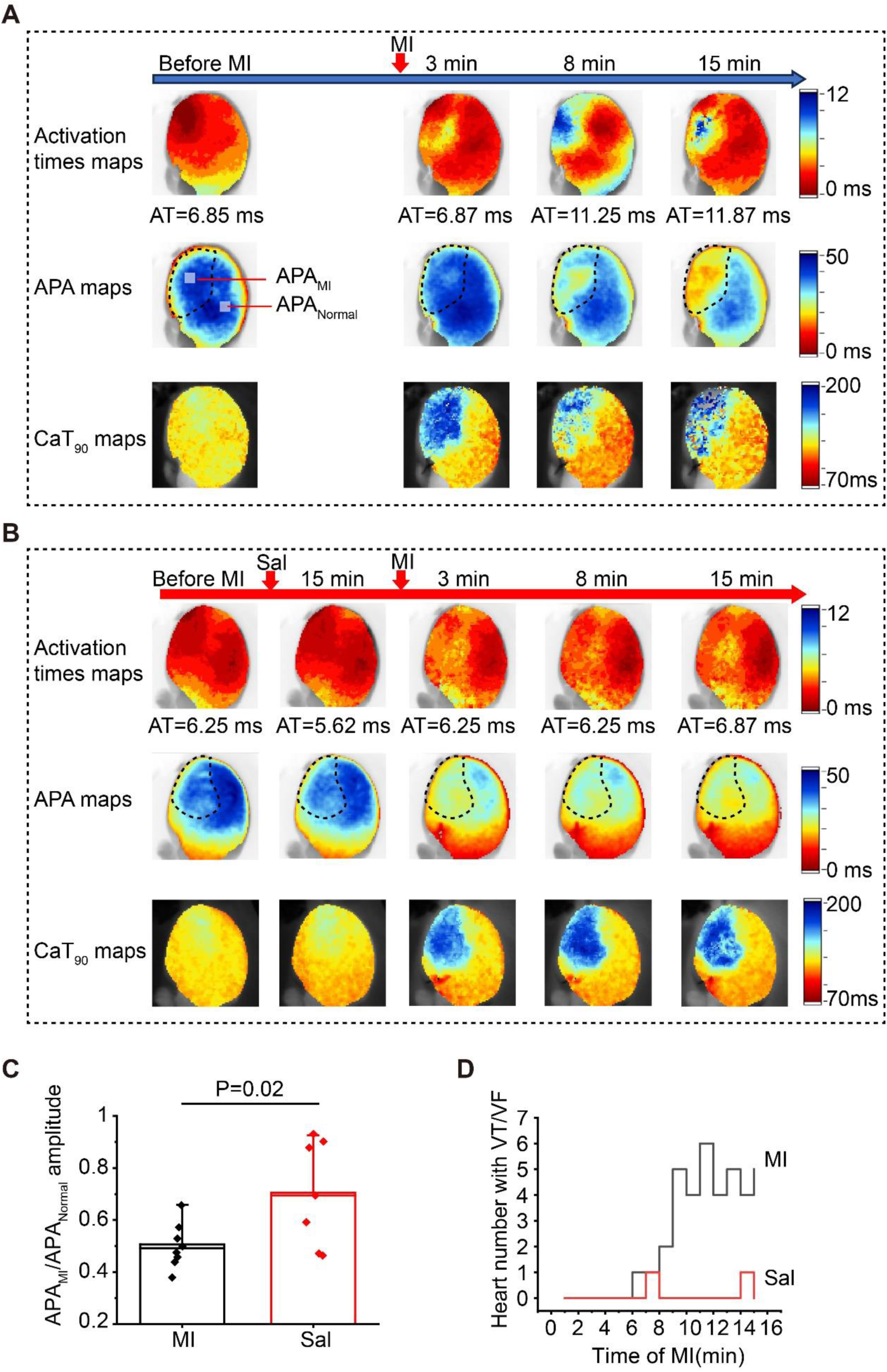
Sal Pre-protection in Myocardial Infarction Models. (A) Electrophysiological changes in isolated rat heart MI models (n=8) before and after LAD ligation. Activation time isochrones, AP amplitude distribution, and calcium transient duration maps show progression from normal conduction to significant conduction delays and reduced AP amplitudes in the infarcted region over time. (B) Effects of Salidroside preconditioning on electrophysiological properties (n=7). Salidroside treatment reduces slow conduction areas, increases AP amplitudes, and confirms myocardial infarction with prolonged calcium transient durations. (C) Ratio of AP amplitude in infarcted regions compared to normal regions. Salidroside preconditioning significantly protects against the reduction in AP amplitude. (D) Incidence of malignant arrhythmias per minute during the ultra-acute phase of MI. Salidroside preconditioning significantly reduces arrhythmia incidence.

Figure 5B displays the activation time isochrones, AP amplitude distribution, and calcium transient duration maps for the Salidroside preconditioning group. After preconditioning with Salidroside and subsequent MI induction, there is a noticeable reduction in slow conduction areas, with increased AP amplitudes in the infarcted region compared to the MI model group. The calcium transient duration maps further confirm the successful induction of myocardial infarction by showing prolonged calcium transient durations in the ischemic regions.

Figure 5C summarizes the ratio of APA in the infarcted region to that in the normal region. Salidroside preconditioning significantly protects against the reduction in AP amplitude caused by ischemia. Additionally, figure 5D shows the number of malignant arrhythmias occurring per minute during the ultra-acute phase (phase IA) of myocardial infarction. Salidroside preconditioning notably reduces the incidence of malignant arrhythmias.

### Effects of Sal Pre-Protection on Conduction Velocity and Conduction Dispersion in Non-Stopbeating Rat Isolated Hearts After Myocardial Infarction

As illustrated in Fig. 6A, we developed an AMI model using ligation of the left anterior descending (LAD) artery. Following ligation, ischemia was observed in the anterior wall of the left ventricle and the apex of the heart. Electrical mapping was performed on the normal region (right ventricle) and the ischemic region (left ventricle) using Pen MEA 1 and Pen MEA 2, respectively. Two 64-channel Pen MEAs were utilized for recording, and pseudo-ECG recordings were synchronized with these measurements. In the control group, premature ventricular contractions (PVCs) were observed after acute myocardial infarction, as indicated by the red arrow in Fig. 6A. The pseudo-ECG analysis revealed a significant elevation in the T wave and widening of the QRS complexes post-MI, whereas the QRS complexes were relatively narrower in the Sal pre-protected group (Fig. 6B), suggesting improved ventricular conduction consistency due to pre-protection.

**Figure 6.**
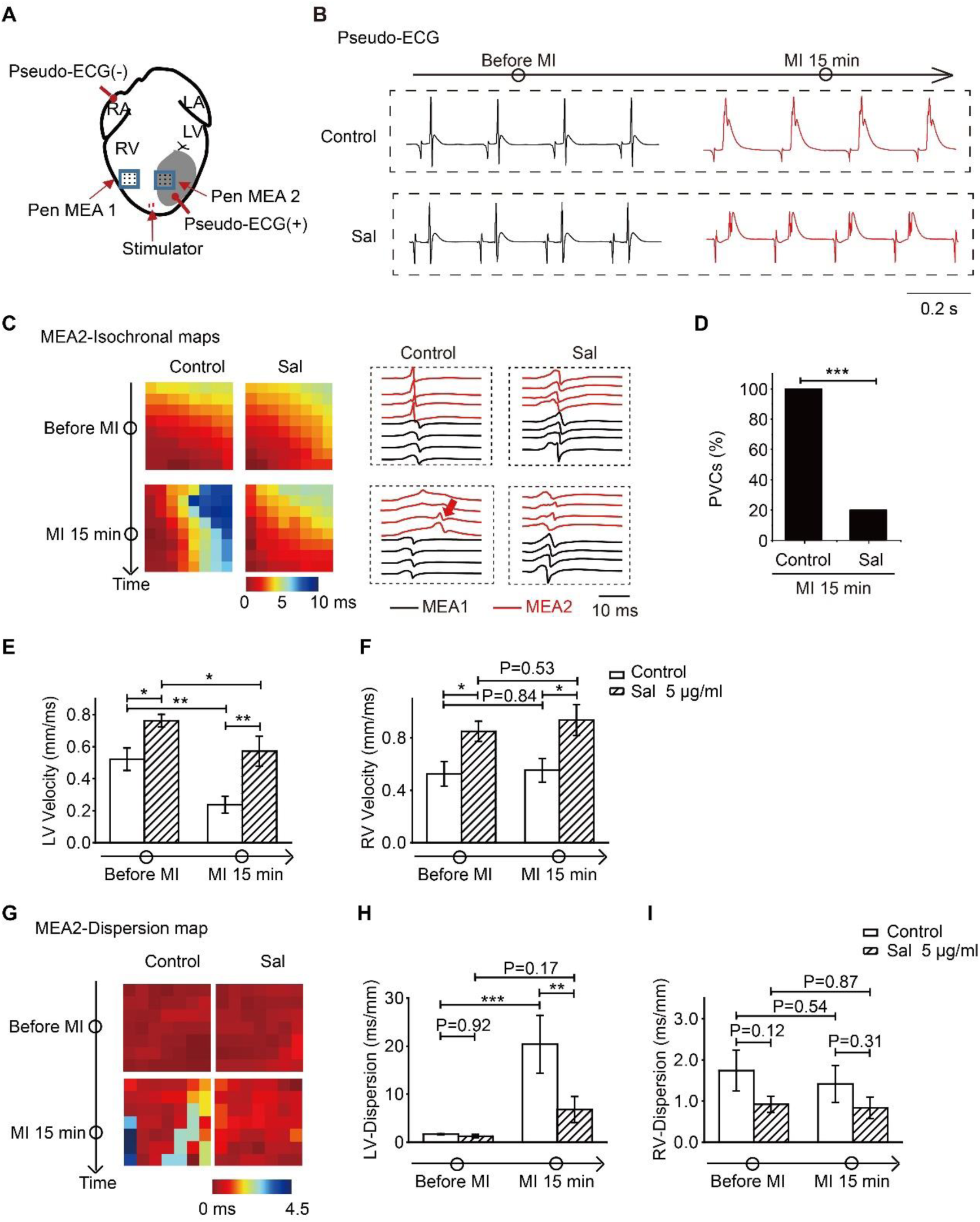
Effects of Sal Pre-protection on Conduction Velocity and Dispersion. (A) AMI model using LAD artery ligation. Electrical mapping and pseudo-ECG recordings from the ischemic and normal regions show PVCs in the control group. (B) Pseudo-ECG analysis post-MI showing elevated T waves and widened QRS complexes in the control group, with improved ventricular conduction in the Sal pre-protected group, n=5-7. (C) Conduction maps from Pen MEA 2 showing uniform conduction direction with cardiac apex stimulation. (D) Incidence of PVCs post-MI. Sal pre-protection significantly reduces PVC occurrence. (E) Conduction velocity in the infarcted area, showing improved conduction with Sal pre-protection. (F) Conduction velocity in the right ventricle before and after MI, with no significant difference between groups. (G) Conduction dispersion analysis showing increased dispersion post-MI in the model group and less dispersion in the Sal pre-protected group. (H) Statistical data on conduction dispersion in the left ventricle under 6 Hz stimulation, showing reduced dispersion in the Sal pre-protected group compared to the model group. (I) Conduction dispersion in the right ventricle paced at 6 Hz, with no significant difference between the Sal group and the AMI model group. *p<0.05, **p<0.01, ***p<0.001.

To ensure uniform conduction direction, cardiac apex stimulation at 6 Hz was employed as shown in Fig. 6C. Representative conduction maps from Pen MEA 2 for both the model group and the Sal pre-protected group before and after myocardial infarction for 15 minutes are depicted. In the control group, PVCs occurred in 100% of the cases (8/8) after acute myocardial infarction, whereas in the Sal pre-protected group, the incidence of PVCs was reduced to 20% (2/10) (Fig. 6D). Statistical analysis indicated that conduction velocity in the myocardial infarction area was significantly reduced after MI. Furthermore, pre-administration of Sal significantly improved conduction velocity (Fig. 6E), with the Sal group demonstrating faster conduction compared to the control group. However, no significant difference was observed in the conduction velocity of the right ventricle before and after MI (Fig. 6F).

Conduction dispersion, which reflects the homogeneity of conduction velocity, was further analyzed. Increased dispersion is a key factor for the occurrence of reentry. Fig. 6G shows conduction dispersion before and after MI in both the AMI group and the Sal pre-protected group. The model group exhibited more significant dispersion, while the Sal group showed less dispersion. Fig. 6H illustrates that before ligation, there was no significant difference in the dispersion of left ventricular conduction between the model group and the Sal group under 6 Hz stimulation (*p* = 0.92). After ligation, dispersion increased in both groups, but the Sal group exhibited a smaller increase compared to the model group (*p* < 0.01). The dispersion in the model group was significantly higher post-ligation compared to pre-ligation (*p* < 0.001). Although the dispersion in the Sal group increased after ligation, the change was not statistically significant compared to pre-ligation. Fig. 6I presents the statistical data for conduction dispersion in the right ventricle paced at 6 Hz. No significant difference was found between the AMI model and the Sal group. In the Sal group, conduction dispersion before and after ligation did not differ significantly from that observed in the AMI group.

### Sal’s Effects on Rabbit Myocardial Infarction

Fig. 7A shows imaging of the rabbit heart during optical mapping. The entire ventral view of the heart was recorded, with pseudo-ECG signals synchronized and obtained from two leads. The stimulation electrode was positioned at the apex of the left ventricle. Fig. 7B illustrates the electrocardiographic changes during myocardial infarction (MI). In the MI model group, ligation caused an immediate inversion of the T wave, followed by a gradual elevation of the ST segment. In contrast, after 15 minutes of Sal pre-protection, the T wave showed mild inversion at 5 minutes, and the ST segment elevation was less pronounced at 25 minutes. Fig. 7C depicts the QRS duration statistics. The QRS duration significantly prolonged with increasing infarction time in the MI model group. However, in the Sal pre-protection group, there was no significant widening of the QRS duration following MI. Fig. 7D shows the location of the stimulation electrode and the pacing frequency set at 4 Hz. The stimulation captured sinus rhythm, with excitation initially occurring at the apex and then propagating gradually along the myocardial wall towards the base. After infarction, the presence of slow conduction areas led to a progressively prolonged activation time. In contrast, the Sal pre-protected hearts did not exhibit noticeable slow conduction areas.

**Figure 7.**
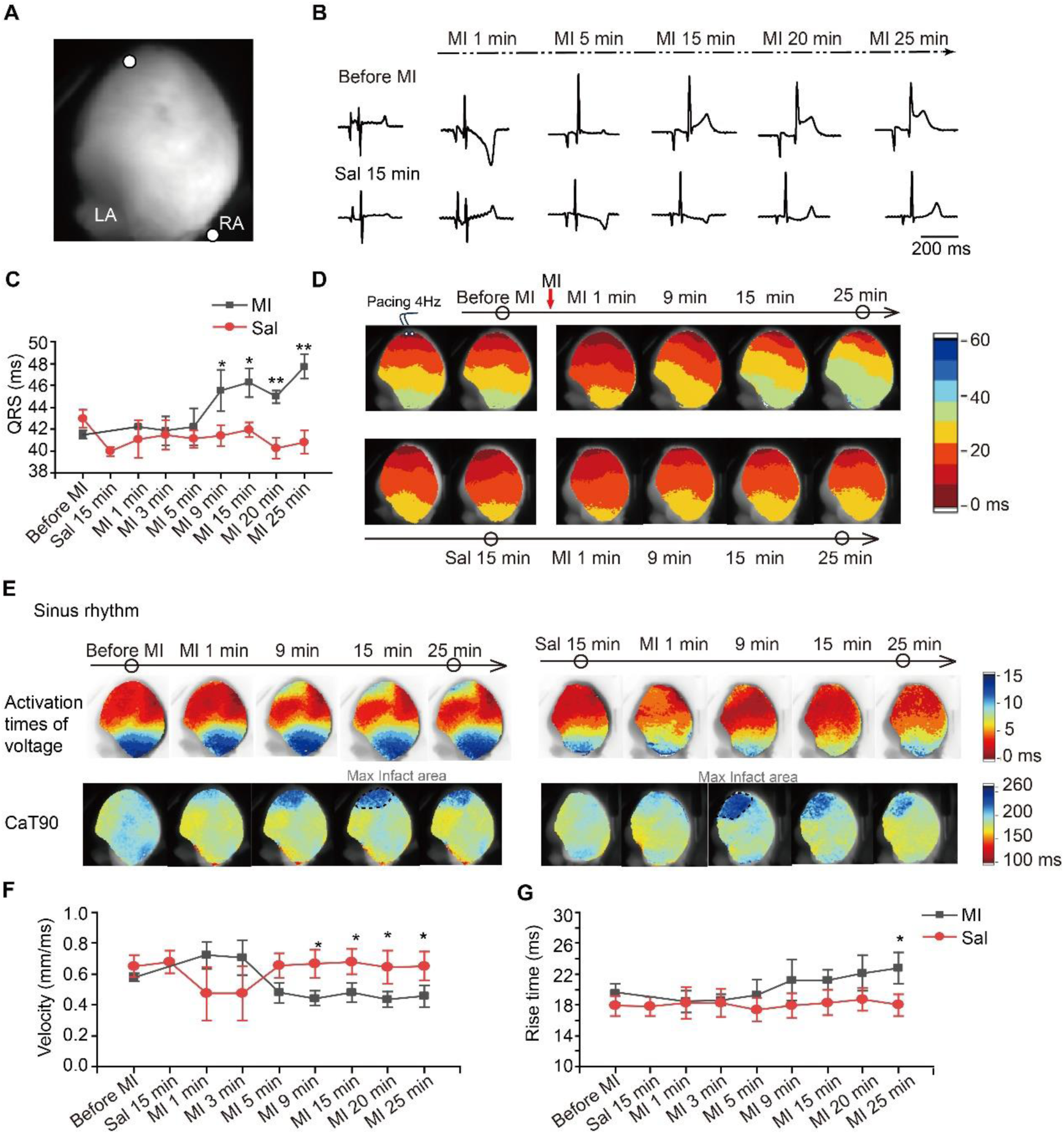
Sal’s Effects on Rabbit Myocardial Infarction. (A) Optical mapping of the rabbit heart during MI, with synchronized pseudo-ECG signals showing stimulation electrode placement. (B) Electrocardiographic changes during MI, with T wave inversion and ST segment elevation in the MI model group (n=4) and milder changes in the Sal pre-protection group (n=5).(C) QRS duration statistics showing prolonged QRS duration in the MI model group, with no significant widening in the Sal pre-protection group.(D) Stimulation electrode placement and pacing frequency in the rabbit heart, with observed slow conduction areas in the MI model group and normal conduction in the Sal pre-protection group.(E) Calcium transient duration maps showing significant conduction delay areas in the MI model group and no distinct delays in the Sal pre-protection group.(F) Conduction velocities and (G) action potential depolarization times, with Sal providing protective effects against depolarization issues caused by hypoxia. *p<0.05, **p<0.01, ***p<0.001.

**Figure 8.**
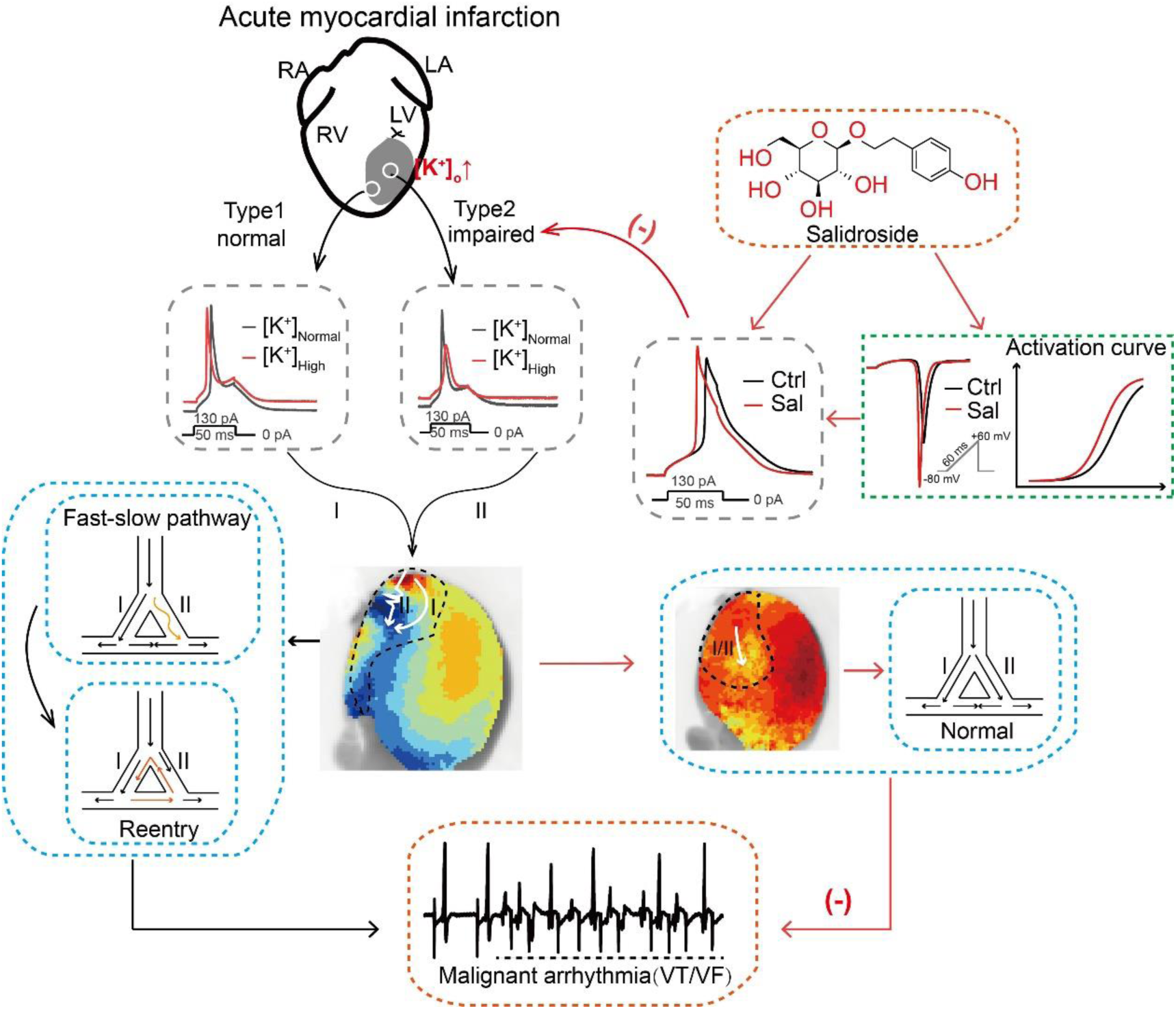
Conceptual Model of Electrophysiological Changes During the Ultra-Acute Phase of Myocardial Infarction (MI) and the Effect of Sal. This diagram illustrates the electrophysiological alterations occurring during the ultra-acute phase of MI and the therapeutic impact of Sal. As MI progresses, extracellular potassium (K⁺) concentration increases, leading to distinct changes within the infarcted myocardial tissue. In the infarction’s central region, severe ischemic damage results in significant electrophysiological impairment. These damaged cells exhibit suppressed excitability due to elevated K⁺, functioning as slow conduction pathways. Conversely, the peripheral areas of the infarcted region experience less damage and maintain relatively normal electrical function. In this high-K⁺ environment, these less affected regions demonstrate increased excitability, potentially leading to ectopic pacing and faster conduction, which can form fast conduction pathways. The coexistence of fast and slow conduction pathways creates favorable conditions for reentry circuits and malignant arrhythmias. Our study proposes that Salidroside (Sal) mitigates these adverse effects by enhancing sodium channel currents and shifting the Nav1.5 activation curve. This modulation facilitates earlier action potential activation and increased amplitude, promoting electrophysiological recovery of damaged myocardial cells. Consequently, Sal reduces the severity of slow conduction pathways and prevents the formation of fast-slow conduction pathways, offering protective effects against arrhythmias during the ultra-acute phase of MI.

Under sinus rhythm, similar to rats, calcium transient duration was used to identify ischemic and hypoxic regions. The first row of Fig. 7E shows that the MI model group exhibited significant conduction delay areas. However, in the Sal pre-protection group, there were no distinct conduction delay regions observed during the MI procedure. Fig. 7F presents the statistical data on conduction velocities for both groups, while Fig. 7G shows the statistics on action potential depolarization times. Both the conduction velocity statistics and depolarization time analysis indicate that Sal provides protective effects against depolarization issues caused by hypoxia.

## Discussion

To summarize our findings, we have developed a conceptual model to explain the electrophysiological changes observed during the ultra-acute phase of myocardial infarction (MI). As the infarction progresses, the concentration of extracellular potassium gradually increases, creating distinct effects within the infarcted area. In the central region of the infarction, myocardial cells experience severe ischemic damage, leading to significant electrophysiological impairment. These damaged cells are more prone to excitability suppression due to the elevated extracellular potassium, making them function as slow conduction pathways. In contrast, the peripheral areas surrounding the infarction are less affected and maintain relatively normal electrical function. In this high-potassium environment, these less damaged regions show increased excitability, which can lead to ectopic pacing and faster conduction, potentially becoming sites for fast conduction pathways. The combination of fast and slow conduction pathways creates conditions conducive to reentry circuits and malignant arrhythmias.

Our study suggests that Salidroside (Sal) can play a crucial role in counteracting these effects. By enhancing sodium channel currents and shifting the Nav1.5 activation curve, Sal facilitates earlier activation of action potentials and increases their amplitude. This action supports the electrophysiological recovery of damaged myocardial cells, reducing the severity of slow conduction pathways. As a result, Sal helps prevent the formation of the fast-slow conduction pathways, thereby offering protection against arrhythmias during the ultra-acute phase of MI.

### Precise Identification of Ischemic Regions and Excitation Heterogeneity

In the study, we precisely identified the ischemic and hypoxic regions within the field of view by leveraging the sensitivity of calcium transient duration to hypoxia, allowing for real-time monitoring of the extent of infarction^26^. The areas with significant changes in calcium transient duration can be referred to as electrically ischemic regions. In previous studies, despite experimental training and efforts to ligate the same location to achieve consistency in infarct regions, it was challenging to ensure the same extent of infarction due to the variability in coronary artery distribution in each heart^27^. This experiment enables the comparison of the electrical conditions of myocardium with equivalent infarction extent by monitoring calcium transients, providing a better understanding of the specific electrical changes in the ischemic regions during myocardial infarction.

A fundamental electrophysiologic concept of arrhythmogenic mechanisms is the interplay between an initiator and a suitable substrate that facilitates the perpetuation of abnormal rhythms^28^. During the Ia phase of myocardial infarction, some infarcted regions exhibit noticeable conduction slowing^29, 30^, while peri-infarct regions display areas with normal conduction and even ectopic beats. Significant conduction block in peri-infarct regions does not emerge until the Ib phase, which is consistent with previous reports indicating severe gap junction uncoupling during that phase^31^. The Ia phase is characterized by dramatic changes in factors such as extracellular potassium concentration, acidity, hypoxia, calcium overload, resting membrane potential, and KATP channel opening^29^. This brief period of heightened arrhythmia incidence suggests that the primary factors may initially excite and subsequently inhibit myocardial activity.

Our data suggest that the different excitability of myocardial cells affected by high extracellular potassium and various electrophysiological injuries could be a major cause of malignant arrhythmias during the Ia phase. Under aerobic conditions, intracellular K+ concentration is high, while extracellular concentration is low, with passive K+ efflux being balanced by active influx through the Na+/K+ pump^32^. During ischemia, this balance is disrupted, resulting in an accumulation of extracellular K+. Within minutes of myocardial ischemia, extracellular K+ concentration increases, raising the resting membrane potential in the ischemic area. Elevated extracellular potassium levels cause the resting membrane potential to rise, leading to the inactivation of Na+ channels^33^. This inactivation decreases Na+ conductance, reducing the amplitude and slope of phase 0, which in turn slows conduction and alters refractoriness^34^. These changes contribute to the complex electrophysiological alterations observed during the early stages of myocardial ischemia.

### Sal acts on cardiac sodium channels, improving electrical conduction in ischemic regions

Using patch-clamp techniques, we observed that Sal exhibited a rapid response and washout effect on acutely isolated Nav1.5 currents, suggesting that Sal likely exerts its effects extracellularly. If it were intracellular, it would not be washed out quickly. The minimal change in action potential amplitude in normal myocardial cells is likely due to the high expression of myocardial sodium channels in these cells, which maintains their rapid depolarization characteristics. Because of the large number of sodium channels, any increase in function is more noticeable when sodium channel function or quantity is reduced. The same effect observed in stable cell lines further indicates that Sal may act directly on Nav channels. Sal shifts the Nav1.5 activation curve to the left, which is an agonistic effect, allowing the channels to activate at more negative voltages. This could be one reason Sal helps maintain relatively normal conduction function in myocardial cells.

### Othe therapeutic effects of Sal

Sal has demonstrated therapeutic effects in mitigating some of these changes. Notably, Sal appears to have a beneficial impact on calcium overload. Supplementary Figure 1 illustrates that Sal reduces calcium alternans in a rabbit model of myocardial infarction (MI) during the ultra-acute phase. This finding is significant because calcium alternans, characterized by irregular fluctuations in calcium transient duration, is a marker of impaired calcium handling and increased arrhythmogenic risk^35^. By attenuating these fluctuations, Sal helps stabilize calcium dynamics and reduce the likelihood of arrhythmias. However, in a rat model of acute myocardial infarction, calcium transients exhibit alternating patterns both before and after the Ia phase, with progressively worsening alternans (Supplementary Figure 2). This observation suggests that calcium transients may not be the primary cause of malignant arrhythmias during the Ia phase.

## Implications for Antiarrhythmic Therapy

Sal offers a novel approach to antiarrhythmic therapy, contrasting with traditional antiarrhythmic drugs. While established antiarrhythmic agents typically target specific ion channels or modulate autonomic nervous system activity^4, 36–39^, Sal’s mechanism appears to address the underlying electrophysiological disturbances in a more comprehensive manner. Unlike conventional drugs that may exhibit narrow therapeutic windows or limited efficacy in certain arrhythmic conditions, Sal demonstrates a unique ability to restore sodium channel function, enhance conduction velocity, and improve overall myocardial electrical stability. This multifaceted action positions Sal as a promising new addition to the antiarrhythmic arsenal, offering a fresh strategy for managing arrhythmias, particularly in the complex context of myocardial infarction.

The relationship between increased conduction velocity and suppression of reentry circuits is a critical aspect of antiarrhythmic therapy. Sal’s ability to enhance conduction velocity can help mitigate the formation of reentry circuits, which are a primary substrate for malignant arrhythmias. By accelerating conduction through damaged myocardium, Sal reduces the time available for reentry circuits to develop, thereby decreasing the risk of sustained arrhythmias. However, increasing conduction velocity is challenging due to the dense expression of sodium channels in each myocardial cell, which complicates efforts to further enhance conduction in normal tissues. Despite this, our studies suggest that in electrically impaired myocardial cells, restoring conduction function is feasible. Sal’s efficacy in this regard is evidenced by its ability to improve conduction velocity in both ischemic and normal myocardial tissues.

Supplementary Figure 4 illustrates that Sal indeed exhibits a concentration-dependent increase in conduction velocity in normal hearts. This finding underscore Sal’s potential to enhance conduction velocity in a manner that is both dose-responsive and functionally significant. Although achieving this effect in damaged myocardium poses additional challenges, Sal’s impact on conduction dynamics highlights its potential as a therapeutic agent for restoring electrical stability in both normal and infarcted hearts. Overall, Sal’s ability to both accelerate conduction and suppress arrhythmogenic reentry circuits marks a significant advancement in antiarrhythmic treatment, offering a promising avenue for future research and clinical application.

## Limitations and Future Directions

Despite the compelling in vitro and ex vivo evidence, our study is limited by the absence of in vivo data. Future research should include animal models and clinical trials to validate Sal’s efficacy and safety in a physiological context^40^. Additionally, the rapid onset and washout of Sal’s effects suggest an extracellular mechanism of action, which warrants further investigation. Identifying Sal’s binding sites on the Nav1.5 channel through computational modeling and mutagenesis studies could provide deeper insights into its molecular interactions and optimize its therapeutic application.

## Clinical Relevance and Recommendations

Our study underscores the importance of considering extracellular potassium dynamics and sodium channel functionality in the management of acute MI. Clinicians should be aware of the potential benefits of incorporating Sal into treatment protocols, particularly for patients at high risk of malignant arrhythmias. Moreover, caution is advised when using Nav1.5 inhibitors in the acute phase of MI, as they may exacerbate conduction abnormalities and arrhythmia risk.

In conclusion, Salidroside offers a novel and effective approach to mitigating malignant arrhythmias during the ultra-acute phase of myocardial infarction by restoring sodium channel function and reducing conduction heterogeneity. These findings pave the way for further research and clinical application, aiming to improve outcomes for patients experiencing acute cardiac events.

## ETHICS STATEMENT

The animal study was reviewed and approved by Animal Research Ethics Committee of Henan SCOPE Research Institute of Electrophysiology.

## AUTHOR CONTRIBUTIONS

Gongxin Wang and Fei Lin initially conceived the project. Yilin Zhao, Chenchen Zhang, Xuefang Li, Xiuming Dong and Dongxu Li performed surgery, ex vivo electrical mapping and patch clamp. Xiulong Wang and Siyu Sun analyzed the data. Gongxin Wang, Chieh-Ju Lu and Yimei Du drafted the manuscript. Xuefang Li and Huan Li reviewed of the data and provided critical advice throughout the research. Guoliang Hao and Zhigang Chen supervised and provided funding for the project. All authors revised the manuscript.

## References

1. Saito Y, Oyama K, Tsujita K, Yasuda S and Kobayashi Y. Treatment strategies of acute myocardial infarction: updates on revascularization, pharmacological therapy, and beyond. J Cardiol. 2023;81:168–178.

2. Zeppenfeld K, Tfelt-Hansen J, de Riva M, Winkel BG, Behr ER, Blom NA, Charron P, Corrado D, Dagres N, de Chillou C, Eckardt L, Friede T, Haugaa KH, Hocini M, Lambiase PD, Marijon E, Merino JL, Peichl P, Priori SG, Reichlin T, Schulz-Menger J, Sticherling C, Tzeis S, Verstrael A, Volterrani M and Group ESCSD. 2022 ESC Guidelines for the management of patients with ventricular arrhythmias and the prevention of sudden cardiac death. Eur Heart J. 2022;43:3997–4126.

3. Anderson JL and Morrow DA. Acute Myocardial Infarction. N Engl J Med. 2017;376:2053–2064.

4. De Groot JR and Coronel R. Acute ischemia-induced gap junctional uncoupling and arrhythmogenesis. Cardiovasc Res. 2004;62:323–34.

5. Sattler SM, Skibsbye L, Linz D, Lubberding AF, Tfelt-Hansen J and Jespersen T. Ventricular Arrhythmias in First Acute Myocardial Infarction: Epidemiology, Mechanisms, and Interventions in Large Animal Models. Front Cardiovasc Med. 2019;6:158.

6. Cox JL, Daniel TM and Boineau JP. The electrophysiologic time-course of acute myocardial ischemia and the effects of early coronary artery reperfusion. Circulation. 1973;48:971–83.

7. Kolettis TM. Ventricular Arrhythmias During Acute Myocardial Ischemia/Infarction: Mechanisms and Management Cardiac Arrhythmias; 2014: 237–251.

8. Taylor J. 2012 ESC Guidelines on acute myocardial infarction (STEMI). Eur Heart J. 2012;33:2501–2.

9. Caughey MC, Arora S, Qamar A, Chunawala Z, Gupta MD, Gupta P, Vaduganathan M, Pandey A, Dai X, Smith SC, Jr. and Matsushita K. Trends, Management, and Outcomes of Acute Myocardial Infarction Hospitalizations With In-Hospital-Onset Versus Out-of-Hospital Onset: The ARIC Study. J Am Heart Assoc. 2021;10:e018414.

10. Wang FS, Lien WP, Fong TE, Lin JL, Cherng JJ, Chen JH and Chen JJ. Terminal cardiac electrical activity in adults who die without apparent cardiac disease. Am J Cardiol. 1986;58:491–5.

11. Chou CC, Zhou S, Hayashi H, Nihei M, Liu YB, Wen MS, Yeh SJ, Fishbein MC, Weiss JN, Lin SF, Wu D and Chen PS. Remodelling of action potential and intracellular calcium cycling dynamics during subacute myocardial infarction promotes ventricular arrhythmias in Langendorff-perfused rabbit hearts. J Physiol. 2007;580:895–906.

12. Eisner DA, Caldwell JL, Kistamas K and Trafford AW. Calcium and Excitation-Contraction Coupling in the Heart. Circ Res. 2017;121:181–195.

13. Feridooni HA, Dibb KM and Howlett SE. How cardiomyocyte excitation, calcium release and contraction become altered with age. J Mol Cell Cardiol. 2015;83:62–72.

14. Herron TJ, Lee P and Jalife J. Optical imaging of voltage and calcium in cardiac cells & tissues. Circ Res. 2012;110:609–23.

15. Zaman S and Kovoor P. Sudden cardiac death early after myocardial infarction: pathogenesis, risk stratification, and primary prevention. Circulation. 2014;129:2426–35.

16. Shaw RM and Rudy Y. Electrophysiologic effects of acute myocardial ischemia: a theoretical study of altered cell excitability and action potential duration. Cardiovasc Res. 1997;35:256–72.

17. Shaw RM and Rudy Y. Electrophysiologic effects of acute myocardial ischemia. A mechanistic investigation of action potential conduction and conduction failure. Circ Res. 1997;80:124–38.

18. Nabavi SF, Braidy N, Orhan IE, Badiee A, Daglia M and Nabavi SM. Rhodiola rosea L. and Alzheimer’s Disease: From Farm to Pharmacy. Phytother Res. 2016;30:532–9.

19. Pu WL, Zhang MY, Bai RY, Sun LK, Li WH, Yu YL, Zhang Y, Song L, Wang ZX, Peng YF, Shi H, Zhou K and Li TX. Anti-inflammatory effects of Rhodiola rosea L.: A review. Biomed Pharmacother. 2020;121:109552.

20. Tao H, Wu X, Cao J, Peng Y, Wang A, Pei J, Xiao J, Wang S and Wang Y. Rhodiola species: A comprehensive review of traditional use, phytochemistry, pharmacology, toxicity, and clinical study. Med Res Rev. 2019;39:1779–1850.

21. Zhang X, Xie L, Long J, Xie Q, Zheng Y, Liu K and Li X. Salidroside: A review of its recent advances in synthetic pathways and pharmacological properties. Chem Biol Interact. 2021;339:109268.

22. Song D, Zhao M, Feng L, Wang P, Li Y and Li W. Salidroside attenuates acute lung injury via inhibition of inflammatory cytokine production. Biomed Pharmacother. 2021;142:111949.

23. Li L, Yang Y, Zhang H, Du Y, Jiao X, Yu H, Wang Y, Lv Q, Li F, Sun Q and Qin Y. Salidroside Ameliorated Intermittent Hypoxia-Aggravated Endothelial Barrier Disruption and Atherosclerosis via the cAMP/PKA/RhoA Signaling Pathway. Front Pharmacol. 2021;12:723922.

24. Xiong Y, Wang Y, Xiong Y and Teng L. Protective effect of Salidroside on hypoxia-related liver oxidative stress and inflammation via Nrf2 and JAK2/STAT3 signaling pathways. Food Sci Nutr. 2021;9:5060–5069.

25. Zhang B, Wang Y, Li H, Xiong R, Zhao Z, Chu X, Li Q, Sun S and Chen S. Neuroprotective effects of salidroside through PI3K/Akt pathway activation in Alzheimer’s disease models. Drug Des Devel Ther. 2016;10:1335–43.

26. MacGowan GA, Du C, Glonty V, Suhan JP, Koretsky AP and Farkas DL. Rhod-2 based measurements of intracellular calcium in the perfused mouse heart: cellular and subcellular localization and response to positive inotropy. J Biomed Opt. 2001;6:23–30.

27. Cao Y, Redd MA, Fang C, Mizikovsky D, Li X, Macdonald PS, King GF and Palpant NJ. New Drug Targets and Preclinical Modelling Recommendations for Treating Acute Myocardial Infarction. Heart Lung Circ. 2023;32:852–869.

28. Sedlis SP. Mechanisms of ventricular arrhythmias in acute ischemia and reperfusion. Cardiovasc Clin. 1992;22:3–18.

29. Stevenson WG, Sager PT, Natterson PD, Saxon LA, Middlekauff HR and Wiener I. Relation of pace mapping QRS configuration and conduction delay to ventricular tachycardia reentry circuits in human infarct scars. J Am Coll Cardiol. 1995;26:481–8.

30. Brunckhorst CB, Stevenson WG, Soejima K, Maisel WH, Delacretaz E, Friedman PL and Ben-Haim SA. Relationship of slow conduction detected by pace-mapping to ventricular tachycardia re-entry circuit sites after infarction. J Am Coll Cardiol. 2003;41:802–9.

31. Cascio WE, Yang H, Muller-Borer BJ and Johnson TA. Ischemia-induced arrhythmia: the role of connexins, gap junctions, and attendant changes in impulse propagation. J Electrocardiol. 2005;38:55–9.

32. Ferrero JM, Gonzalez-Ascaso A and Matas JFR. The mechanisms of potassium loss in acute myocardial ischemia: New insights from computational simulations. Front Physiol. 2023;14:1074160.

33. Remme CA and Bezzina CR. Sodium channel (dys)function and cardiac arrhythmias. Cardiovasc Ther. 2010;28:287–94.

34. Klabunde RE. Cardiac electrophysiology: normal and ischemic ionic currents and the ECG. Adv Physiol Educ. 2017;41:29–37.

35. Qian YW, Clusin WT, Lin SF, Han J and Sung RJ. Spatial heterogeneity of calcium transient alternans during the early phase of myocardial ischemia in the blood-perfused rabbit heart. Circulation. 2001;104:2082–7.

36. Al-Khatib SM, Stevenson WG, Ackerman MJ, Bryant WJ, Callans DJ, Curtis AB, Deal BJ, Dickfeld T, Field ME, Fonarow GC, Gillis AM, Granger CB, Hammill SC, Hlatky MA, Joglar JA, Kay GN, Matlock DD, Myerburg RJ and Page RL. 2017 AHA/ACC/HRS guideline for management of patients with ventricular arrhythmias and the prevention of sudden cardiac death: Executive summary: A Report of the American College of Cardiology/American Heart Association Task Force on Clinical Practice Guidelines and the Heart Rhythm Society. Heart Rhythm. 2018;15:e190–e252.

37. Amoni M, Dries E, Ingelaere S, Vermoortele D, Roderick HL, Claus P, Willems R and Sipido KR. Ventricular Arrhythmias in Ischemic Cardiomyopathy-New Avenues for Mechanism-Guided Treatment. Cells. 2021;10.

38. Di Diego JM and Antzelevitch C. Ischemic ventricular arrhythmias: experimental models and their clinical relevance. Heart Rhythm. 2011;8:1963–8.

39. Lei M, Wu L, Terrar DA and Huang CL. Modernized Classification of Cardiac Antiarrhythmic Drugs. Circulation. 2018;138:1879–1896.

40. Martinez-Navarro H, Minchole A, Bueno-Orovio A and Rodriguez B. High arrhythmic risk in antero-septal acute myocardial ischemia is explained by increased transmural reentry occurrence. Sci Rep. 2019;9:16803.

